# Genome-wide specificity profiles of CRISPR-Cas Cpf1 nucleases in human cells

**DOI:** 10.1101/057802

**Authors:** Benjamin P. Kleinstiver, Shengdar Q. Tsai, Michelle S. Prew, Nhu T. Nguyen, Moira M. Welch, Jose M. Lopez, Zachary R. McCaw, Martin J. Aryee, J. Keiths Joung

**Affiliations:** Molecular Pathology Unit, Massachusetts General Hospital, Charlestown, MA 02129 USA; Center for Cancer Research, Massachusetts General Hospital, Charlestown, MA 02129 USA; Center for Computational and Integrative Biology, Massachusetts General Hospital, Charlestown, MA 02129 USA; Department of Pathology, Harvard Medical School, Boston, MA 02115 USA; Department of Biostatistics, Harvard T.H. Chan School of Public Health, Boston, Massachusetts 02115, USA

## Abstract

CRISPR-Cas Cpf1 nucleases have recently been described as an alternative genome-editing platform^1^, yet their activities and genome-wide specificities remain largely undefined. Here we show that two Cpf1 nucleases function robustly in human cells with on-target efficiencies comparable to those of the widely used *Streptococcus pyogenes* Cas9 (**SpCas9**)^2–5^. We also demonstrate that four to six bases at the 3’ end of the short CRISPR RNA (**crRNA**) used to program Cpf1 are insensitive to single base mismatches but that many of the other bases within the crRNA targeting region are highly sensitive to single or double substitutions. Consistent with these results, GUIDE-seq and targeted deep sequencing analyses of two Cpf1 nucleases revealed no detectable off-target cleavage for over half of 20 different crRNAs we examined. Our results suggest that the two Cpf1 nucleases we characterized generally possess high specificities in human cells, a finding that should encourage broader use of these genome editing enzymes.

Clustered, regularly interspaced, short palindromic repeat (**CRISPR**) systems encode RNA-guided endonucleases that are essential for bacterial adaptive immunity^6^. Much work has shown that these CRISPR-associated (**Cas**) nucleases can be easily programmed to cleave target DNA sequences of interest for genome editing applications in a variety of different organisms^2–5^. One class of these nucleases, known as Cas9 proteins, naturally complex with two short RNAs: a crRNA and a trans-activating crRNA (**tracrRNA**)^7, 8^SpCas9, the most commonly used Cas9 orthologue, uses a crRNA harboring 20 nucleotides at its 5’ end with complementarity to the “protospacer” region of its target DNA site. Efficient cleavage also requires the presence of a protospacer adjacent motif (**PAM**) which is recognized by SpCas9. The crRNA and tracrRNA are now typically combined into a single ~100 nt guide RNA (**gRNA**)^7, 9–11^, which functions efficiently with SpCas9 to direct its cleavage activity. The genome-wide specificities of SpCas9 nucleases paired with various gRNAs have been well-characterized using a variety of different approaches^12–15^. Engineered SpCas9 variants with substantially improved genome-wide specificities have also recently been described^16, 17^.

Recent classification and characterization of Cpf1 proteins from type V CRISPR-Cas systems has revealed that a Cas protein called Cpf1 can also be programmed to cleave specific target DNA sequences^1, 18–20^. In contrast to SpCas9, Cpf1 requires only a single 42 nt crRNA, which bears 23 nucleotides at its 3’ end that are complementary to the protospacer of a target sequence^1^. Furthermore, whereas SpCas9 recognizes an NGG PAM sequence that lies 3’ to the protospacer, Cpf1 orthologues from both *Acidaminococcus sp. BV3L6* and *Lachnospiraceae bacterium ND2006* (**AsCpfl** and **LbCpfl**, respectively) generally recognize TTTN PAMs that are positioned 5’ to the protospacer^1^. Initial characterization of AsCpf1 and LbCpf1 showed that these nucleases can be programmed to edit target sites in human cells^1^; however, their robustness remains largely undefined because to date these nucleases have only been tested on a small number of endogenous gene sites. In addition, the genome-wide specificities of these two nucleases have not been characterized but remain an important parameter to be defined if AsCpf1 and LbCpf1 are to be broadly used for research and therapeutic applications.

In initial experiments, we assessed the robustness of AsCpf1 and LbCpf1 nucleases in U2OS human cells by testing their activities against 19 target sites for in four different human genes, with each site proximal to a DNA sequence that can be efficiently mutagenized by SpCas9 (**Supplementary Fig. 1a**). Six of these sites (DNMT1 sites 1 – 4 and EMX1 sites 1 and 2) had been previously targeted with Cpf1 nucleases in HEK293 cells^1^. Both AsCpf1 and LbCpf1 efficiently induced insertion or deletion mutations (**indels**) at all 19 of these sites as judged by T7 Endonuclease I (**T7E1**) mismatch assays, with modification percentages ranging from 3.8% to 56.2% (mean of 26.7%) for AsCpf1 and 5.2% to 53.2% (mean of 33.8%) for LbCpf1 (**Fig. 1a**). By comparison, the modification percentages for the ten proximal sites targeted by SpCas9 ranged from 26.8% to 55.4% (mean of 42.5%) (**Supplementary Fig. 1b**).

**Figure 1.**
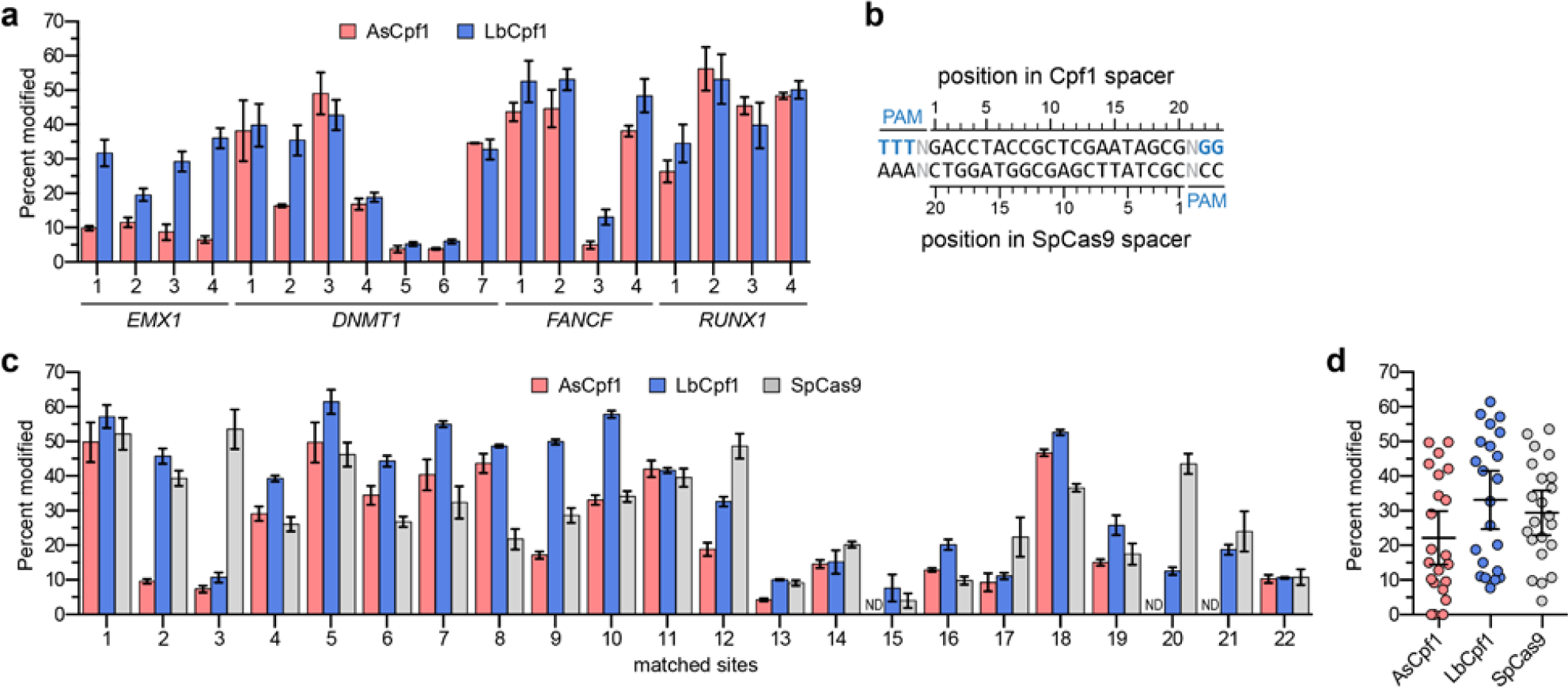
On-target indel mutation percentages induced by Cpf1 nucleases in human cells. (**a**) Endogenous gene percent modification induced by AsCpf1 and LbCpf1 at 19 endogenous human gene sites (determined by T7E1 assay). Target sites were chosen in amplicons known to be efficiently modified by SpCas9 (**Supplementary Fig. 1**). Error bars, s.e.m.; n = 3. (**b**) and (**c**) Matched target sites for AsCpf1, LbCpf1, and SpCas9 that share a common protospacer sequence (**panel b**) were examined for mutagenesis by AsCpf1, LbCpf1, and SpCas9 nucleases (determined by T7E1 assay; **panel c**). Error bars, s.e.m.; n = at least 3; ND, not detected. (**d**) Summary of matched site on-target activities from **panel c**, with means and 95% confidence intervals shown.

To more directly compare the on-target genome editing activities of AsCpf1 and LbCpf1 with that of SpCas9, we identified 22 additional endogenous human gene target sites (**Supplementary Table 1**) that contained overlapping target sites for both classes of nucleases (**Fig. 1b**). These matched sites were selected to have variable numbers of *in silico* Cas-OFFinder^21^ predicted off-target sites in the genome (**Supplementary Table 2**). Analysis of on-target activities on these 22 sites revealed variable modification efficiencies by AsCpf1, LbCpf1, and SpCas9 (**Figs. 1c** and **1d**). Time-course analysis revealed that near-maximum indel mutagenesis was achieved by 72 hours post-transfection (**Supplementary Fig. 2**).

Next, to initially assess the specificity of AsCpf1 and LbCpf1, we examined the tolerance of these nucleases to mismatches at the crRNA-protospacer DNA interface. To accomplish this, we mismatched adjacent pairs of bases within the complementarity regions of crRNAs targeted to three different endogenous target sites in the human *DNMT1* gene (**Fig. 2a**). With all three target sites, we found that two adjacent mismatched bases at positions 1 through 18 (with 1 being the most PAM-proximal base) virtually eliminated detectable site modification as assessed by T7E1 assay for both AsCpf1 and LbCpf1 (**Fig. 2a**). By contrast, two adjacent mismatches of the PAM distal bases between positions 19 and 23 had substantially less pronounced effects on the activities of both Cpf1 nucleases (**Fig. 2a**). Further testing of AsCpf1 and LbCpf1 at two of the three *DNMT1* target sites using crRNAs bearing single mismatches positioned along the length of the protospacer complementarity region revealed substantial and consistent tolerance to mismatches at positions 1, 8, 9, and 19 through 23 (**Fig. 2b**). Taken together, these experiments suggested that both AsCpf1 and LbCpf1 can be highly sensitive to mismatched crRNA nucleotides at most positions between 1 and 18, and that recognition of bases at the 3’ end of the crRNA may be dispensable for efficient nuclease activity.

**Figure 2.**
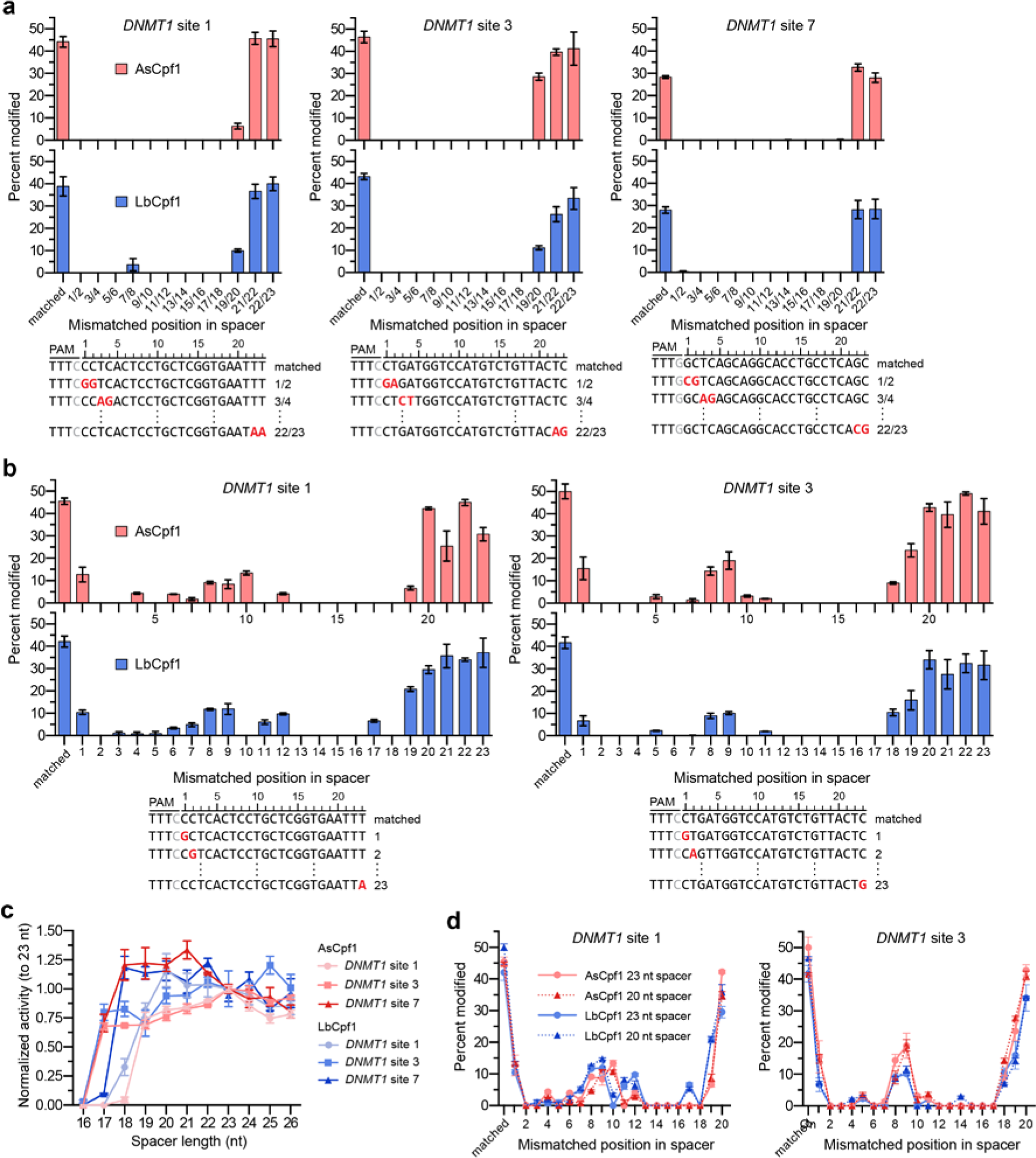
Tolerance of AsCpf1 and LbCpf1 to mismatched or truncated crRNAs. (**a**, **b**) Endogenous gene modification by AsCpf1 and LbCpf1 using crRNAs that contain pairs of mismatched bases (panel **a**) or singly mismatched bases (panel **b**). Activity determined by T7E1 assay; error bars, s.e.m.; n = 3. (**c**) Summary of mutagenesis percentages by AsCpf1 and LbCpf1 at 3 different endogenous sites with crRNAs bearing variable length 3’ end truncations or extensions (see also **Supplementary Fig. 3a**), where activities are normalized to mutagenesis observed when using the canonical 23 nucleotide spacer. Error bars, s.e.m.; n = 3; nt, nucleotide. (**d**) Activity of AsCpf1 and LbCpf1 when programed with singly mismatched crRNAs either of 23 or 20 nucleotides in length. For clarity, data from **panel b** is re-presented here. Activity determined by T7E1 assay; error bars, s.e.m.; n = 3.

We further tested the importance of bases at the 3’ end of the crRNA by evaluating the activities of AsCpf1 and LbCpf1 with a series of crRNAs bearing variable-length 3’ end deletions and extensions. Examination of Cpf1 activity when paired with truncated crRNAs against three *DNMT1* sites revealed that mutagenesis efficiencies remained robust even when four to six bases were removed from the 3’ end of the crRNA (**Fig. 2c**; **Supplementary Fig. 3a**). Interestingly, the addition of up to three matched bases to the 3’ end of the canonical length 23 nt crRNAs did not substantially alter indel formation by either AsCpf1 or LbCpf1 (**Fig. 2c** and **Supplementary Fig. 3a**). Collectively, our findings suggest that only 17 to 19 bases of crRNA protospacer complementarity are required for robust Cpf1 activity and that four to six bases at the 3’ end of a 23 nt spacer may not be necessary for efficient target site cleavage. Previous studies with SpCas9 showed that gRNAs bearing truncated target complementarity regions are generally more sensitive to single mismatches at the gRNA/target DNA interface than their full-length counterparts^22^. However, we observed virtually no differences in AsCpf1 and LbCpf1 activities when using singly mismatched crRNAs bearing 20 instead of 23 nucleotide spacer sequences (**Fig. 2d**; **Supplementary Fig. 3b**).

To profile the genome-wide specificities of Cpf1 nucleases in human cells, we used the Genome-wide Unbiased Identification of Double-stranded breaks Enabled by sequencing (GUIDE-seq) method^12^. GUIDE-seq has been previously used by our group and others to identify off-target cleavage sites of wild-type and engineered Cas9 nucleases^12, 16, 23–27^based on the capture of an end-protected, double-stranded oligodeoxynucleotide (dsODN) into nuclease-induced breaks in living cells with subsequent amplification and identification of adjacent genomic sequences by deep sequencing. We assessed dsODN incorporation at 11 of the 22 matched sites we had shown above can be targeted by all three nucleases (**Fig. 1c**) and observed efficient tag integration with AsCpf1, LbCpf1, and SpCas9 at all of these genomic loci in human U2OS cells as judged by RFLP assay (**Supplementary Fig. 4a**). The overall on-target modification efficiencies observed in these experiments (the combination of dsODN integration and indels) were also determined using the T7EI mismatch assay (**Supplementary Fig. 4b**).

The ratios of dsODN incorporation efficiency to overall modification efficiency observed with AsCpf1 and LbCpf1 at these 11 sites, a measure of the efficiency of tag integration at the on-target sites, were lower than the ratios observed with SpCas9 (means of 0.31, 0.28, and 0.53 for AsCpf1, LbCpf1, and SpCas9, respectively; **Supplementary Figs. 4c** and **4d**). We also assessed dsODN tag integration (**Supplementary Fig. 4e**) with AsCpf1 and LbCpf1 at 9 of the 19 human gene target sites we used in the experiments of **Fig. 1a** and again found reasonably efficient tag incorporation and on-target modification in U2OS cells (**Supplementary Fig. 4f**). Ratios of dsODN incorporation efficiency to overall modification efficiency for AsCpf1 and LbCpf1 were comparable with these nine crRNAs (**Supplementary Fig. 4g**). Collectively, these results demonstrate that dsODN tag integration occurs at Cpf1-induced breaks, albeit with somewhat lower efficiencies than what is observed for SpCas9, perhaps due to differences into the nature of the DSBs induced by these nucleases (blunt ends for SpCas9 and 5’ overhangs for Cpf1; ref 1). Variability in the location of dsODN tag integration was higher than typically observed with SpCas9, but generally appeared to be constrained to 10-20 bps surrounding the expected DSB position (**Fig. 3a**).

**Figure 3.**
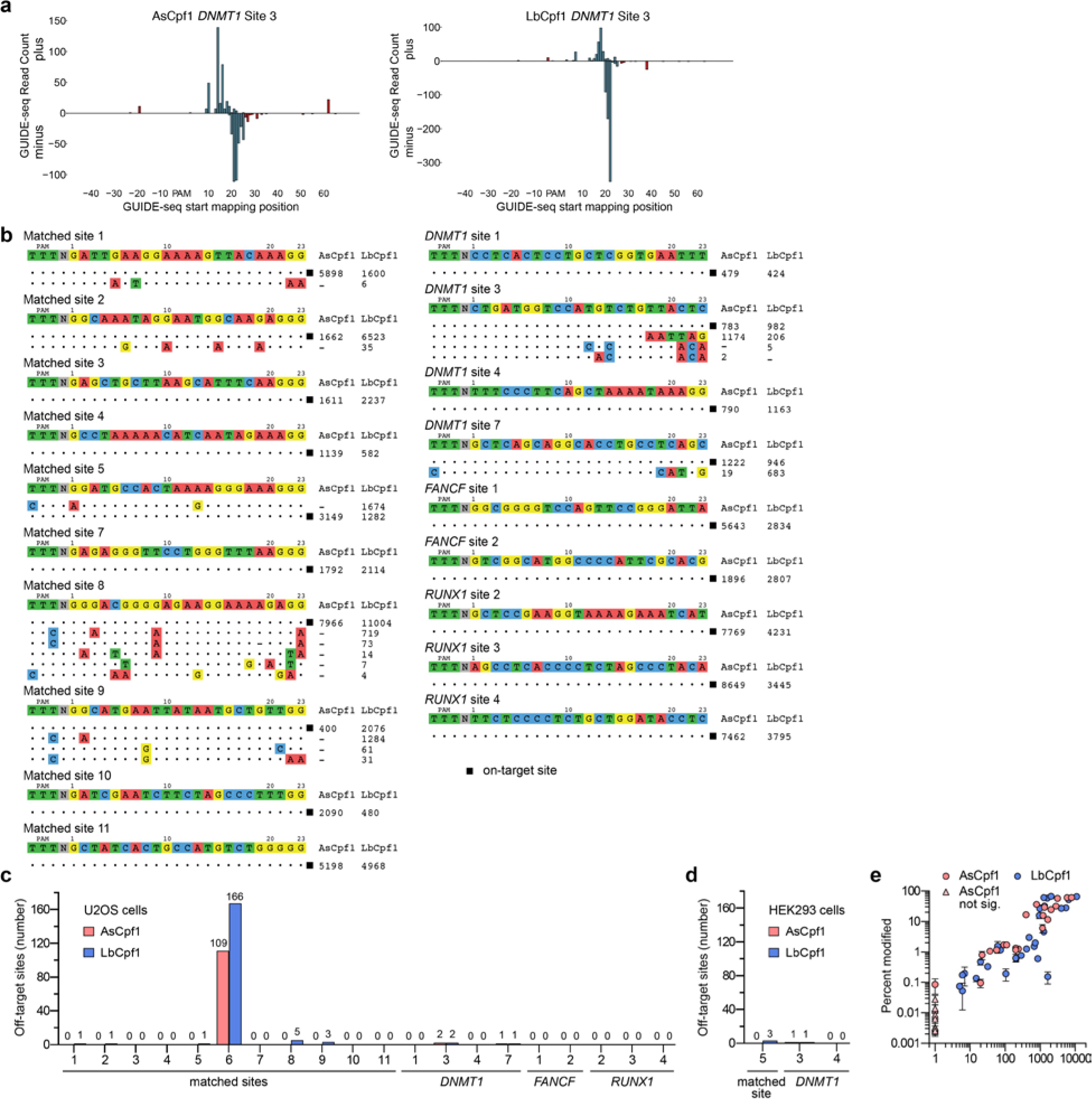
Genome-wide specificities of AsCpf1 and LbCpf1. (**a**) Relative GUIDE-seq read start mapping position. The first 5’ base of the protospacer adjacent to the PAM is position 1. Reads that start within the protospacer region are colored in blue, reads that originate outside are colored in red. Reads mapping to the plus strand are depicted above the x-axis, reads mapping in the reverse orientation are depicted below. (**b**) Off-target sites for AsCpf1 and LbCpf1 with 20 different crRNAs, determined using GUIDE-seq in U2OS cells. Mismatched positions in thetarget sites of off-targets are highlighted in color, and GUIDE-seq read counts shown to the right of the on- and off-target sequences represent a measure of cleavage efficiency at a given site. (**c**) and (**d**) Summary of the total number of off-target sites detected by GUIDE-seq in U2OS cells with 20 crRNAs (**panel c**) and in HEK293 cells with 3 crRNAs (**panel d**). The off-target sequences and GUIDE-seq read counts for matched site #6 are shown in **Supplementary Fig. 5**. (**e**) Scatterplots of mean mutagenesis versus GUIDE-seq read counts analyzed from independent samples. Mutagenesis percentages for AsCpf1 that are not significantly different from negative controls are indicated with triangles. Error bars, s.e.m.; n = 3.

Having established reasonably efficient tag incorporation at Cpf1-induced DNA breaks for 20 different crRNAs, we performed GUIDE-seq using these crRNAs and both Cpf1 nucleases in U2OS cells (a total of 40 GUIDE-seq experiments). As expected, the on-target sites for each of the 20 crRNAs tested with AsCpf1 and LbCpf1 were readily identified by GUIDE-seq (**Fig. 3b**). Strikingly, no crRNA-specific off-target sites were detected by GUIDE-seq for 17 of the 20 crRNAs with AsCpf1 and for 12 of the 20 crRNAs with LbCpf1 (**Figs. 3b** and **3c**; **Supplementary Table 4**). The remaining crRNAs induced cleavage at only a very small number of off-target sites (range of 1 to 5 per crRNA), with the exception of the crRNA for matched site 6 for which 109 and 166 off-target sites were identified for AsCpf1 and LbCpf1, respectively (**Figs. 3b** and **3c**; **Supplementary Fig. 5**). The larger numbers of off-target cleavage sites identified for the matched site 6 crRNA may be due to the relatively greater number of closely matched potential off-target sequences (as predicted by Cas-OFFinder^21^) that occur in the human genome for this crRNA as compared with the others (**Supplementary Table 2**). Some of the off-target sites we identified possess multiple mismatched bases at the PAM-distal end of the protospacer, consistent with the results of our experiments with systematically mismatched crRNAs. AsCpf1 generally targeted fewer numbers of off-target sites than LbCpf1 in matched comparisons and also appeared to more stringently specify the PAM, consistent with previous *in vitro* experiments showing that AsCpf1 is more specific for a TTTN PAM than LbCpf1 (ref 1).

To extend our findings to another cell line, we performed GUIDE-seq experiments with AsCpf1 and LbCpf1 in human HEK293 cells using three of the 20 crRNAs we had tested in U2OS cells. RFLP and T7E1 experiments showed GUIDE-seq tag incorporation and overall mutagenesis rates to be slightly lower in HEK293 cells compared with U2OS cells (**Supplementary Figs. 6a-6c**), consistent with previous studies^26^. GUIDE-seq analysis with each of the three crRNAs and AsCpf1 or LbCpf1 in HEK293 cells identified a similar number of off-target sites to what was observed in corresponding experiments performed in U2OS cells (**Figs. 3c** and **3d**). Although some off-target sites were the same in both cell types, some were also unique to one cell type and showed some differences in GUIDE-seq read count numbers (**Supplementary Fig. 6d**). Of note, the one crRNA that did not identify any off-target sites with both AsCpf1 and LbCpf1 in U2OS cells also failed to identify any sites in HEK293 cells (**Supplementary Fig. 6d**).

Given the less efficient dsODN incorporation we observed with Cpf1 nucleases relative to Cas9, we sought to assess the sensitivity of GUIDE-seq for detecting Cpf1-induced off-target mutations. As previously described for Cas9, we did this by performing targeted amplicon sequencing to quantify mutagenesis at off-target sites (detected by GUIDE-seq) in U2OS cells transfected without the dsODN. We sequenced 30 GUIDE-seq-identified off-target sites induced by LbCpf1 with eight different crRNAs, and also sequenced these same sites for AsCpf1 even though 10 were not identified by GUIDE-seq. Mutation percentages and GUIDE-seq read counts correlated well and revealed that GUIDE-seq can detect off-target sites mutagenized in the ~0.1% range (**Fig. 3e** and **Supplementary Table 3**). Importantly, for the AsCpf1-induced off-target sites not identified by GUIDE-seq, we observed mutation percentages that were either not statistically significant and/or at levels below 0.01% (**Fig. 3e** and **Supplementary Table 3**).

Because we found relatively few Cpf1-induced off-target sites using GUIDE-seq, we examined whether the method was detecting indel mutations occurring at closely mismatched sites in the human genome. To do this, we performed targeted amplicon sequencing of 61 of the most closely mismatched potential off-target sites for 15 of the 20 crRNAs we had assessed by GUIDE-seq. For 51 of these 61 mismatched sites, we failed to find significant evidence of indels in U2OS cells transfected with Cpf1 nuclease and a crRNA but no dsODN tag (**Supplementary Fig. 7**). Among the ten mismatched sites at which we observed significant indel mutagenesis above background (0.026% to 32.0%), six had been found by our GUIDE-seq experiments and the other four not identified by GUIDE-seq had very low levels of mutations (0.026% to 0.206%) (**Supplementary Fig. 7** and **Supplementary Table 3**). Based on these data, we conclude that Cpf1 nucleases generally do not appear to efficiently mutagenize the majority of closely related sequences and that GUIDE-seq can detect Cpf1 off-target sites that are mutagenized with frequencies as low as the ~0.1−0.2% range.

*In silico* analysis demonstrates that Cpf1 nucleases with a TTTN PAM have at least 1 target site in a high percentage of first- and second-coding exons, microRNAs, and promoter regions in human cells, including some regions that cannot be targeted with wild-type SpCas9 (although this expansion in targeting range diminishes with removal of the 5’ G required when using an RNA polymerase III U6 promoter to express gRNAs for SpCas9; **Supplementary Fig. 8**).

Our results establish that AsCpf1 and LbCpf1 nucleases can robustly induce indel mutations at endogenous gene target sites in human cells with efficiencies comparable to what can be achieved with wild-type SpCas9 nuclease. GUIDE-seq analysis and targeted deep sequencing using a large number of different crRNAs with both AsCpf1 and LbCpf1 suggests that these nucleases are generally highly specific in human cells, with the majority of nucleases showing no evidence of off-target effects detectable by GUIDE-seq. Furthermore, the high sensitivity of Cpf1 to single base mismatches in certain positions of the protospacer may enable these nucleases to be used for allele-specific editing of heterozygous alleles. Although it is challenging, (if not impossible) to directly compare the genome-wide specificities of Cas9 and Cpf1 nucleases due to their different modes of recognition and PAM requirements, our analysis suggests that the specificities of Cpf1 nucleases (and of AsCpf1 in particular) may approach that of recently described high-fidelity SpCas9 variants^16, 17^. Improving the detection limit of GUIDE-seq will enable better assessment of whether lower frequency off-target effects are induced by these nucleases. In addition, it may be possible to further improve Cpf1 nuclease specificities using strategies similar to those recently applied to SpCas9^16, 17^. Collectively, our study should encourage broader use of Cpf1 for research applications and exploration of the use of these nucleases for the development of highly specific therapeutics.

## METHODS

### Plasmids and oligonucleotides

A list and partial sequences of plasmids used in this study can be found in **Supplementary Note 1**; crRNA sequences are listed in **Supplementary Table 1** and oligonucleotide sequences are found in **Supplementary Table 5**. AsCpf1 and LbCpf1 human expression plasmids (SQT1659 and SQT1665, respectively) were generated by inserting the open-reading frames of these nucleases from plasmids pY010 and pY016 (Addgene plasmids # 69982 and # 69988, respectively) into the NotI and AgeI sites of pCAG-CFP (Addgene # 11179). Plasmid SQT817 was used to express SpCas9 (ref ^28^). Oligonucleotide duplexes corresponding to spacer sequences were annealed and ligated into BsmBI-digested BPK3079, BPK3082, and BPK1520 (ref ^24^) for U6 promoter-driven expression of AsCpf1, LbCpf1, and SpCas9 gRNAs, respectively. New plasmids described in this study will be deposited with the non-profit plasmid repository Addgene: http://www.addgene.org/crispr-cas.

### Human cell culture and transfection

U2OS cells (obtained from Toni Cathomen, Freiburg) and HEK293 cells were cultured at 37°C with 5% CO_2_ in Advanced DMEM supplemented with 10% heat-inactivated fetal bovine serum, 2 mM GlutaMax, and penicillin/streptomycin (all cell culture products were obtained from Life Technologies). Cell line identity was validated by STR profiling (ATCC) and mycoplasma testing was performed twice per month. 2 × 10^5^ cells were transfected using the SE Cell Line Nucleofector Kit with 500 ng Cpf1- or SpCas9-encoding plasmid and 250 ng gRNA expression plasmid and the DN-100 program for U2OS cells, or with 300 ng Cpf1- or SpCas9-encoding plasmid and 150 ng gRNA expression plasmid using the CM-137 program for HEK293 cells, on a 4D-Nucleofector according to the manufacturer’s instructions (Lonza). For negative control transfections, Cpf1- or SpCas9-encoding plasmids were co-transfected with a U6-null plasmid that did not encode a crRNA. Genomic DNA extraction was performed approximately 72 hours following nucleofection using an Agencourt DNAdvance Genomic DNA Isolation Kit (Beckman Coulter).

### T7E1 assays

T7 endonuclease I (T7E1) mutation detection assays were performed as previously described^29^ to determine Cpf1 or SpCas9 editing efficiencies at endogenous loci in human cells. Briefly, target loci were PCR amplified using Phusion Hot Start Flex DNA Polymerase (New England Biolabs) with 100 ng of genomic DNA as template. Primers used for PCR amplification are listed in **Supplementary Table 5**. Following an Agencourt AMPure XP (Beckman Coulter) purification, 200 ng of PCR product was denatured, reannealed, and digested with T7E1 (New England Biolabs). AMPure XP purified digestion reactions were analyzed using a QIAxcel capillary electrophoresis instrument (Qiagen), enabling calculation of estimated modification percentages.

### GUIDE-seq

GUIDE-seq experiments were performed essentially as previously described^12^. Briefly, U2OS or HEK293 cells were transfected as described above with the addition of either 100 or 5 pmol, respectively, of an end-protected double-stranded oligodeoxynucleotide (dsODN) GUIDE-seq tag that encodes an internal NdeI restriction site. Restriction-fragment length polymorphism (RFLP) assays, performed as previously described^24^, and T7E1 assays were performed at the target loci to determine tag integration efficiencies and modification percentages, respectively. High-throughput sequencing libraries generated after tag-specific amplification were sequenced using an Illumina MiSeq sequencer as previously described. Data was analyzed using open-source *guide-seq* software^30^ with a window size of 75 bp and allowing for up to 7 mismatches relative to the intended target site. The threshold of 7 mismatches was chosen as the value where less than 0.001% of alignments would be expected to occur by chance (one locus excluded by this threshold showed medium to high mapping read counts in all 40 GUIDE-seq experiments (i.e., all 20 crRNAs with AsCpf1 or LbCpf1; **Supplementary Table 4**). This segment mapped to the U6 promoter that is present within chromosome 15 region but more detailed sequence analysis revealed that these reads are derived from the U6 promoter present on the crRNA expression vector used in our GUIDE-seq experiments (data not shown). These reads likely result from ligation of dsODN to the presence of some linearized crRNA vector present in our plasmid preparations). High-confidence cell-type specific SNPs were called using SAMTools. Updated *guide-seq* software to analyze Cpf1 GUIDE-seq experiments is available online at: http://www.jounglab.org/guideseq/.

### Targeted deep sequencing experiments

Targeted deep sequencing experiments were performed as previously described^16^. On-target, GUIDE-seq identified off-target, and potential off-target loci predicted using Cas-OFFinder^21^ were PCR amplified using Phusion Hot Start Flex DNA Polymerase (New England Biolabs) with the primers listed in **Supplementary Table 5**. A high-throughput library preparation kit (KAPA BioSystems) was used to generate dual-indexed Tru-seq libraries that were sequenced on an Illumina MiSeq sequencer. Amplicons with less than 5000 mapping read counts were excluded from the analysis. P-values were calculated by fitting a negative binomial regression model comparing indel rates between nuclease treated and control samples. To avoid model fitting issues when all control samples report zero reads, 1 count was added to each member of the set of observed read counts. Deep sequencing data and the results of statistical tests are
reported in **Supplementary Table 3**.

### Targeting range calculations

Targetable sites for SpCas9 and Cpf1 were identified using Cas-OFFinder^21^ by searching for genomic sequences matching GN_19_-NGG/N_20_-NGG and TTTN-N_23_ motifs, respectively. Targeting range calculations are based on overlaps between predicted cut sites and features defined in the RefSeq gene table downloaded from the UCSC Genome Browser on January 23, 2016. The first (defined as the most 5’) and second exons were extracted for each unique gene name. The set of TSS upstream sequences (200bp and 1000bp) were defined using all transcripts, and only counted once in cases where two genes share the same TSS upstream sequence. The set of microRNAs was defined as those transcripts with a “MIR” prefix in their RefSeq gene names.

## Acknowledgments

This work was supported by a National Institutes of Health (NIH) Director’s Pioneer Award (DP1 GM105378) and NIH R01 GM107427 to J.K.J., the Jim and Ann Orr Research Scholar Award (to J.K.J.), a Natural Sciences and Engineering Research Council of Canada Postdoctoral Fellowship (to B.P.K.), and an MGH Tosteson Award (to S.Q.T.). New reagents described in this work will be deposited with the non-profit plasmid distribution service Addgene (http://www.addgene.org/crispr-cas).

## Author Contributions

B.P.K., S.Q.T., and J.K.J. conceived of and designed experiments. B.P.K., S.Q.T., M.S.P., N.T.N., and M.M.W. performed all human cell experiments and analyzed data. S.Q.T., J.M.L., Z.M., and M.J.A. analyzed the GUIDE-seq and targeted deep-sequencing data. B.P.K., S.Q.T., and J.K.J. wrote the manuscript with input from all authors.

## References

1. Zetsche, B. et al. Cpf1 Is a Single RNA-Guided Endonuclease of a Class 2 CRISPR-Cas System. Cell 163, 759–771 (2015).

2. Sander, J.D. & Joung, J.K. CRISPR-Cas systems for editing, regulating and targeting genomes. Nat Biotechnol 32, 347–355 (2014).

3. Hsu, P.D., Lander, E.S. & Zhang, F. Development and applications of CRISPR-Cas9 for genome engineering. Cell 157, 1262–1278 (2014).

4. Doudna, J.A. & Charpentier, E. Genome editing. The new frontier of genome engineering with CRISPR-Cas9. Science 346, 1258096 (2014).

5. Maeder, M.L. & Gersbach, C.A. Genome-editing Technologies for Gene and Cell Therapy. Mol Ther (2016).

6. Wright, A.V., Nunez, J.K. & Doudna, J.A. Biology and Applications of CRISPR Systems: Harnessing Nature’s Toolbox for Genome Engineering. Cell 164, 29–44 (2016).

7. Jinek, M. et al. A programmable dual-RNA-guided DNA endonuclease in adaptive bacterial immunity. Science 337, 816–821 (2012).

8. Deltcheva, E. et al. CRISPR RNA maturation by trans-encoded small RNA and host factor RNase III. Nature 471, 602–607 (2011).

9. Cong, L. et al. Multiplex genome engineering using CRISPR/Cas systems. Science 339, 819–823 (2013).

10. Mali, P. et al. RNA-guided human genome engineering via Cas9. Science 339, 823–826 (2013).

11. Jinek, M. et al. RNA-programmed genome editing in human cells. Elife 2, e00471 (2013).

12. Tsai, S.Q. et al. GUIDE-seq enables genome-wide profiling of off-target cleavage by CRISPR-Cas nucleases. Nat Biotechnol 33, 187–197 (2015).

13. Frock, R.L. et al. Genome-wide detection of DNA double-stranded breaks induced by engineered nucleases. Nat Biotechnol 33, 179–186 (2015).

14. Wang, X. et al. Unbiased detection of off-target cleavage by CRISPR-Cas9 and TALENs using integrase-defective lentiviral vectors. Nat Biotechnol 33, 175–178 (2015).

15. Kim, D. et al. Digenome-seq: genome-wide profiling of CRISPR-Cas9 off-target effects in human cells. Nat Methods 12, 237–243, 231 p following 243 (2015).

16. Kleinstiver, B.P. et al. High-fidelity CRISPR-Cas9 nucleases with no detectable genome-wide off-target effects. Nature 529, 490–495 (2016).

17. Slaymaker, I.M. et al. Rationally engineered Cas9 nucleases with improved specificity. Science 351, 84–88 (2016).

18. Schunder, E., Rydzewski, K., Grunow, R. & Heuner, K. First indication for a functional CRISPR/Cas system in Francisella tularensis. Int J Med Microbiol 303, 51–60 (2013).

19. Makarova, K.S. et al. An updated evolutionary classification of CRISPR-Cas systems. Nat Rev Microbiol 13, 722–736 (2015).

20. Fagerlund, R.D., Staals, R.H. & Fineran, P.C. The Cpf1 CRISPR-Cas protein expands genome-editing tools. Genome Biol 16, 251 (2015).

21. Bae, S., Park, J. & Kim, J.S. Cas-OFFinder: a fast and versatile algorithm that searches for potential off-target sites of Cas9 RNA-guided endonucleases. Bioinformatics 30, 1473–1475 (2014).

22. Fu, Y., Sander, J.D., Reyon, D., Cascio, V.M. & Joung, J.K. Improving CRISPR-Cas nuclease specificity using truncated guide RNAs. Nat Biotechnol 32, 279–284 (2014).

23. Kleinstiver, B.P. et al. Broadening the targeting range of Staphylococcus aureus CRISPR-Cas9 by modifying PAM recognition. Nat Biotechnol (2015).

24. Kleinstiver, B.P. et al. Engineered CRISPR-Cas9 nucleases with altered specificities. Nature 523, 481–485 (2015).

25. Yin, H. et al. Therapeutic genome editing by combined viral and non-viral delivery of CRISPR system components in vivo. Nat Biotechnol (2016).

26. Bolukbasi, M.F. et al. DNA-binding-domain fusions enhance the targeting range and precision of Cas9. Nat Methods (2015).

27. Friedland, A.E. et al. Characterization of Staphylococcus aureus Cas9: a smaller Cas9 for all-in-one adeno-associated virus delivery and paired nickase applications. Genome Biol 16, 257 (2015).

28. Tsai, S.Q. et al. Dimeric CRISPR RNA-guided FokI nucleases for highly specific genome editing. Nat Biotechnol 32, 569–576 (2014).

29. Reyon, D. et al. FLASH assembly of TALENs for high-throughput genome editing. Nat Biotechnol 30, 460–465 (2012).

30. Tsai, S.Q., Topkar, V.V., Joung, J.K. & Aryee, M.J. guideseq: open-source software for analysis of GUIDE-seq data. Nat Biotech in press (2016).

